# Ridge and crossrib height of butterfly wing scales is a toolbox for structural color diversity

**DOI:** 10.1101/2024.03.28.585318

**Authors:** Cédric Finet, Qifeng Ruan, Yi Yang Bei, Vinodkumar Saranathan, Antónia Monteiro

**Affiliations:** Biological Sciences, National University of Singapore, Singapore 117543, Singapore; Ministry of Industry and Information Technology Key Lab of Micro-Nano Optoelectronic Information System, Harbin Institute of Technology, Shenzhen 518055, P. R. China; Guangdong Provincial Key Laboratory of Semiconductor Optoelectronic Materials and Intelligent Photonic Systems, Harbin Institute of Technology, Shenzhen 518055, P. R. China; Division of Science, Yale-NUS College, National University of Singapore, 138609, Singapore; NUS Nanoscience and Nanotechnology Initiative (NUSNNI), National University of Singapore, 117581, Singapore; Division of Sciences, School of Interwoven Arts and Sciences, Krea University, Sricity, India

**Keywords:** Biophotonics, structural coloration, butterfly scale, ridge, crossrib, interference

## Abstract

The brightest and most vivid colors of butterflies usually originate from light reflecting off the cuticular scales that cover the wing membrane. These scales have an intricate architecture that consists of an upper layer, a grid of longitudinal ridges and transverse crossribs, connected to a lower lamina by pillars called trabeculae. Whereas the role of the lower lamina as a reflector has been well documented in simpler scales, this study unveils the role of the scales’ upper surface in generating or fine-tuning hue, brightness, and saturation. In the nymphalid *Bicyclus anynana*, we showed that changes in ridge and trabecula heights accompanied changes in hue of scales produced via artificial selection. We further found that this correlation between ridge height and hue can be generalized to 40 scale types from 35 species across butterfly families. By combining focused ion beam milling, microspectrophotometry, and optical modelling, we found that modifying the ridge height is sufficient to change ridge hue, notably in *Morpho didius* whose blue color was thought to be generated exclusively by lamella protruding from ridges, rather than ridge height. This study identifies the scale’s upper surface as a toolbox for structural color diversity in butterflies and proposes a geometrical model to predict color that unifies species with and without *Morpho*-type Christmas-tree ridges.

## INTRODUCTION

Organismal structural colors result from the reflection of specific wavelengths of light by sub-micrometer structures found in biological materials ^1–3^. In butterflies, structural colors originate primarily from the scales that cover the wing membrane ^4–6^. These scales, around 100 µm in length, are the dead cuticular skeletons of single epidermal cells that grow and project out of the epithelial layer during pupal development. Scales are highly diverse in morphology ^7–9^, but most share a common Bauplan with an upper surface made of a grid of longitudinal ridges and transverse crossribs defining open windows. Pillars, or trabeculae, support this grid structure on a lower lamina of finite thickness ^10^. This Bauplan is believed to represent the scales found in the last common ancestor of extant butterflies ^11^.

The current paradigm for structural color production in these architecturally simple, stereotypical scales is via light interacting with the lower lamina that acts as a thin film reflector ^12, 13^. Variation in the thickness of this lower lamina across nymphalid butterflies changes the color hue produced by thin-film interference ^13–16^. Similar thin-film structural coloration has been described in scales formed of a fused upper and lower lamina in extant butterflies ^17, 18^ and primitive Lepidoptera ^19, 20^. This current simple model, however, is incomplete as it does not consider the contribution of the upper lamina (in non-fused scales) for color generation. Here we demonstrate the important role of this upper surface, specifically of the longitudinal ridges, in generating structural coloration even in stereotypical “simple” butterfly wing scales.

## RESULTS

### The upper surface substantially contributes to overall scale coloration

To disentangle the relative contributions of upper surface and lower lamina on scale coloration, we measured and compared how light was reflected from the upperside (abwing side) and underside (adwing side) surfaces of 40 scales with stereotypical morphologies from 35 species (Figure 1A). This allowed us to quantify how much the upper surface modulates the thin-film structural color produced by the lower lamina. We restricted our sampling to scales with the simple, “ancestor-like” Bauplan. Thus, we excluded broadband reflective scales that contain a contiguous upper lamina with no windows ^21–25^, as well as scales having a modified lumen filled with multilayers or highly-ordered photonic crystals ^10, 26–29^.

**Figure 1.**
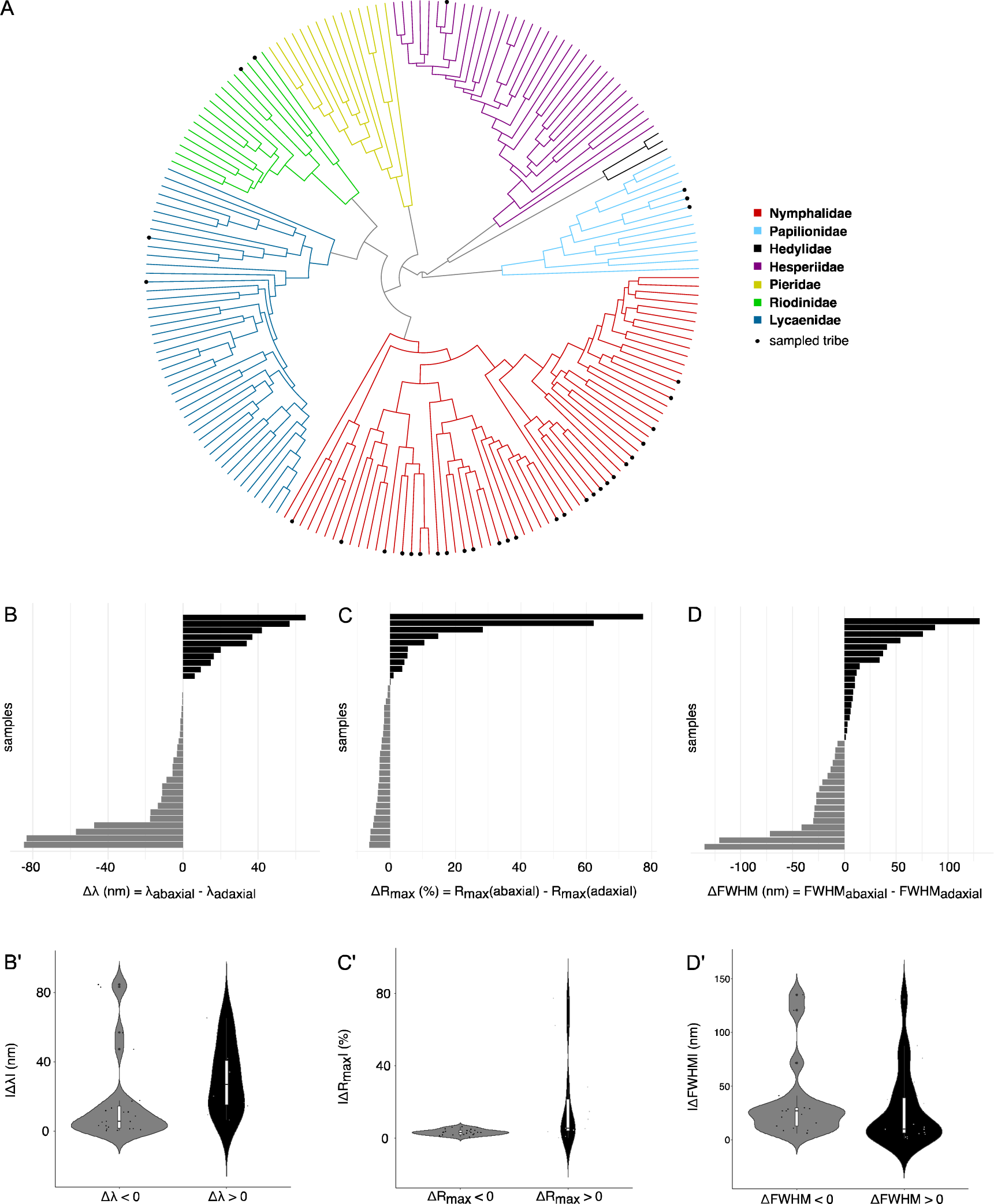
Contribution of upper surface to overall scale coloration. (A) Samples used in this study are mapped onto the phylogeny of butterflies by Espeland *et al.* 2018 ^63^. (B-D) Shift in peak reflectance, reflectance intensity, and saturation between upper surface and lower lamina across sample scales. (B’-D’) Distribution of the absolute value of shift in peak reflectance, reflectance intensity, and saturation, towards lower or higher wavelengths in the visible light spectrum. The central line in the violin plot indicates the median of the distribution, while the top and bottom of the box represent the third and first quartiles of the data, respectively. The whiskers show up to 1.5 times the inter-quartile range.

We first investigated how the presence of the upper surface impacted hue, reflectance intensity, and saturation in our sample of 40 scales. We found that the presence of an upper surface nearly always shifted the reflectance peak, or hue, of the lower lamina to either higher or lower wavelengths depending on the sample (Figure 1B). The shift in reflectance peak ranged from approximately 0 to 85 nm, without a significant directional bias (Figure 1B’). The upper surface could also either increase or decrease the reflectance intensity of a scale (Figure 1C). In the majority of species, the upper surface decreased reflectance by a small amount, but in a small proportion of species, this surface contributed to a very large increase in reflectance intensity (Figure 1C’). For instance, the intensity of the blue ground scales in *Morpho sulkowskyi* increases by 77% through light reflecting from its upper surface nanostructures. Finally, we found that the upper surface can also either increase or decrease color “purity”, or saturation, similarly in a bi-directional manner (Figure 1D, D’). In summary, the upper surface — and its associated nanostructures — impacts the final hue and brightness of the scale in most of the scales investigated in this study.

### Artificial selection acts on the geometry of the upper surface to evolve structural color

To further examine the contribution of the upper surface of scales in the generation of color, we revisited an artificial selection experiment performed on scale hue in *Bicyclus anynana* butterflies. This experiment produced violet-blue scales from UV-reflecting brown ground scales over eight generations of artificial selection in the laboratory. The overall violet-blue coloration was shown to result, at least in part, from an increased thickness of the lower lamina ^14^. By re-examining conserved, dried specimens used to establish the violet-blue lines, we now further observed that the ridges and crossribs also acquired a violet-blue coloration (Figures 2A, A’). We quantified this change in color by measuring the reflectance of the ridges and crossribs on upper surfaces manually isolated from the underlying lower lamina.

**Figure 2.**
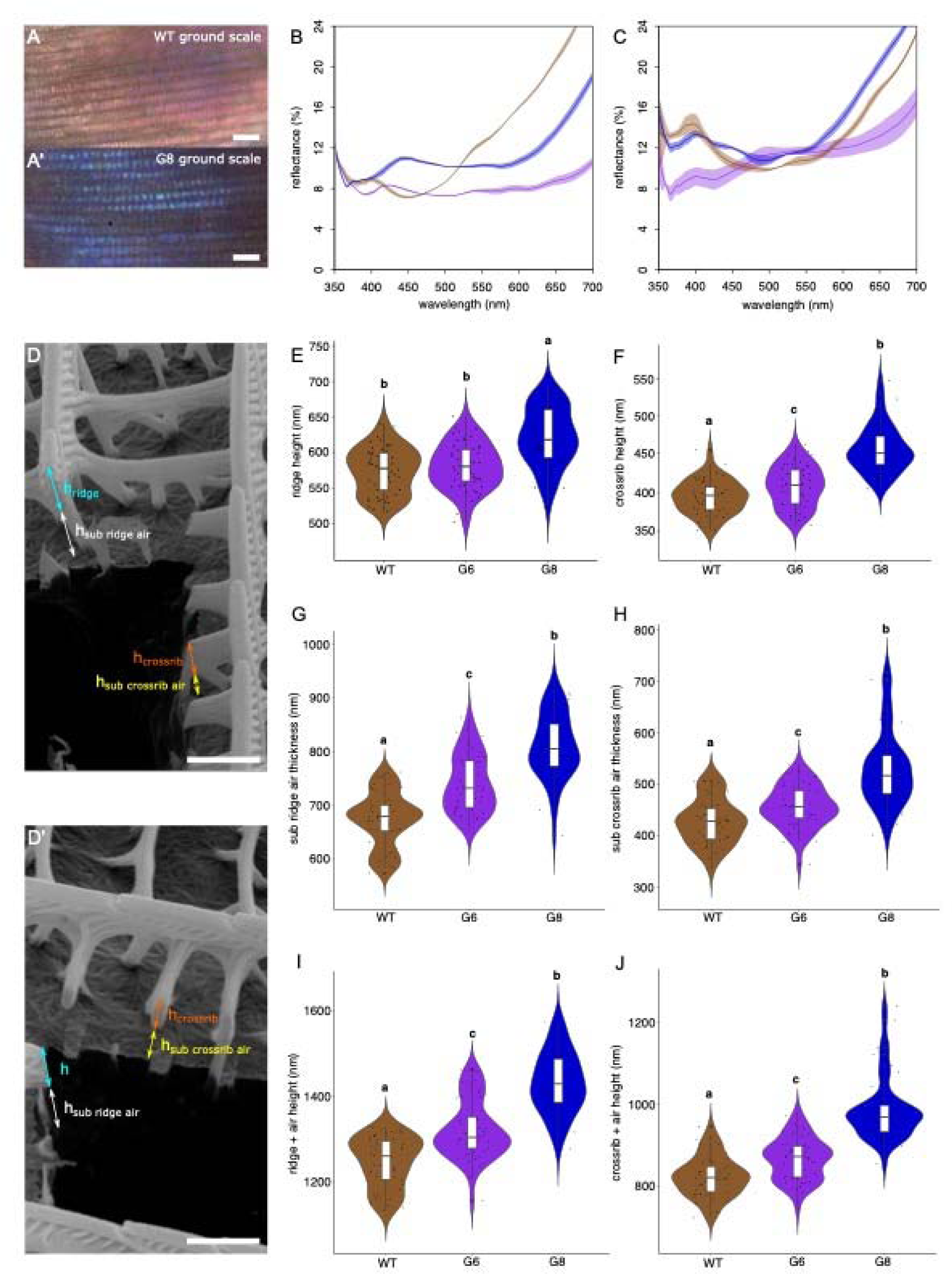
Evolution of scale reflectance and thicknesses of upper surface nanostructures over the course of an artificial selection experiment in *B. anynana* that targeted (blue) scale hue. (A, A’) Optical microscopy images of the abwing side of WT and violet-blue line (G8) ground scales. Scale bar is 4 μm. (B, C) Measured reflectance spectra of ridges and crossribs over generations. Graph colors were arbitrarily chosen: WT scales (brown), violet-blue line (G6) scales (purple), violet-blue line (G8) scales (blue). (D, D’) SEM images of the abwing side of G8 ground scales after FIB milling, showing both transverse (D) and sagittal (D’) views. (E, F) Ridge and crossrib height over generations. (G, H) Air layer thickness under ridge and crossrib over generations. (I, J) Total height over generations. Means sharing the same letter are not significantly different (Tukey-adjusted comparisons). The central line in the violin plot indicates the median of the distribution, while the top and bottom of the box represent the third and first quartiles of the data, respectively. The whiskers show up to 1.5 times the inter-quartile range.

We found that the reflectance peak for the ridges and crossribs shifted towards blue wavelengths over the course of the experiment (Figure 2B, C). By measuring the height of ridges and crossribs from transverse and sagittal focused ion beam scanning electron micrograph (FIB-SEM) sections (Figure 2D, D’), we also found that the height of both structures increased over the course of the experiment (Figure 2E, F). Similar trends were found for the thicknesses of the air layer below the ridges (Figure 2G) and crossribs (Figure 2H), as well as for the total height (chitin + air layer) of ridges (Figure 2I) and crossribs (Figure 2J). Lastly, we detected an increase in the mean distance between crossribs (d_WT_ = 862±71 nm versus d_G8_ = 1160±116 nm) (Figure S1B) (Tables S2 and S3), but not between ridges (Figure S1C). The increase in the distance between crossribs increased the mean window area (Figure S1D) (area_WT_ = 0.72±0.2 μm^2^ versus area_G8_ = 1.2±0.3 μm^2^) (Tables S2 and S3). These results suggest that artificial selection acted simultaneously on several components of the scale geometry: the lower lamina thickness as previously identified ^14^, but also the ridge and crossrib heights, as well as the spacing between crossribs (this study). These morphological changes all contribute to the color shift observed over several generations of artificial selection in *B. anynana*.

### Scale ridge hue correlates with ridge height

To test if conclusions drawn from the *B. anynana* selection lines can be generalized to other species, we measured both ridge reflectance and geometries in the same 40 stereotypical scale types from the 35 species sampled above, but using intact scales. We first investigated how the reflectance measurements on ridges made in manually separated scales of *B. anynana* compared to those made in intact scales of the G8 generation (Figures S2A and S2B). The wavelength of peak light reflection was identical regardless of sampling strategy (Figure S2C). Furthermore, reflectance measurements centered on the ridges and on the lower lamina had distinct wavelength peaks (Figure S2C), demonstrating that our microspectrophotometer enables the measurement of individual ridge reflectance. Measurements of ridge reflectance of all subsequent species were, therefore, done on intact scales. Under epi-illumination, ridges exhibited a continuum of colors ranging from violet to green (Figures 3A-F). The morphology and the geometry of these colored ridges (and crossribs) varied substantially across scales (Figures 3A’-F’ and 3A’’-F’’), including cases where ridges and/or crossribs had the appearance of chitinous walls with almost no underlying air gaps.

**Figure 3.**
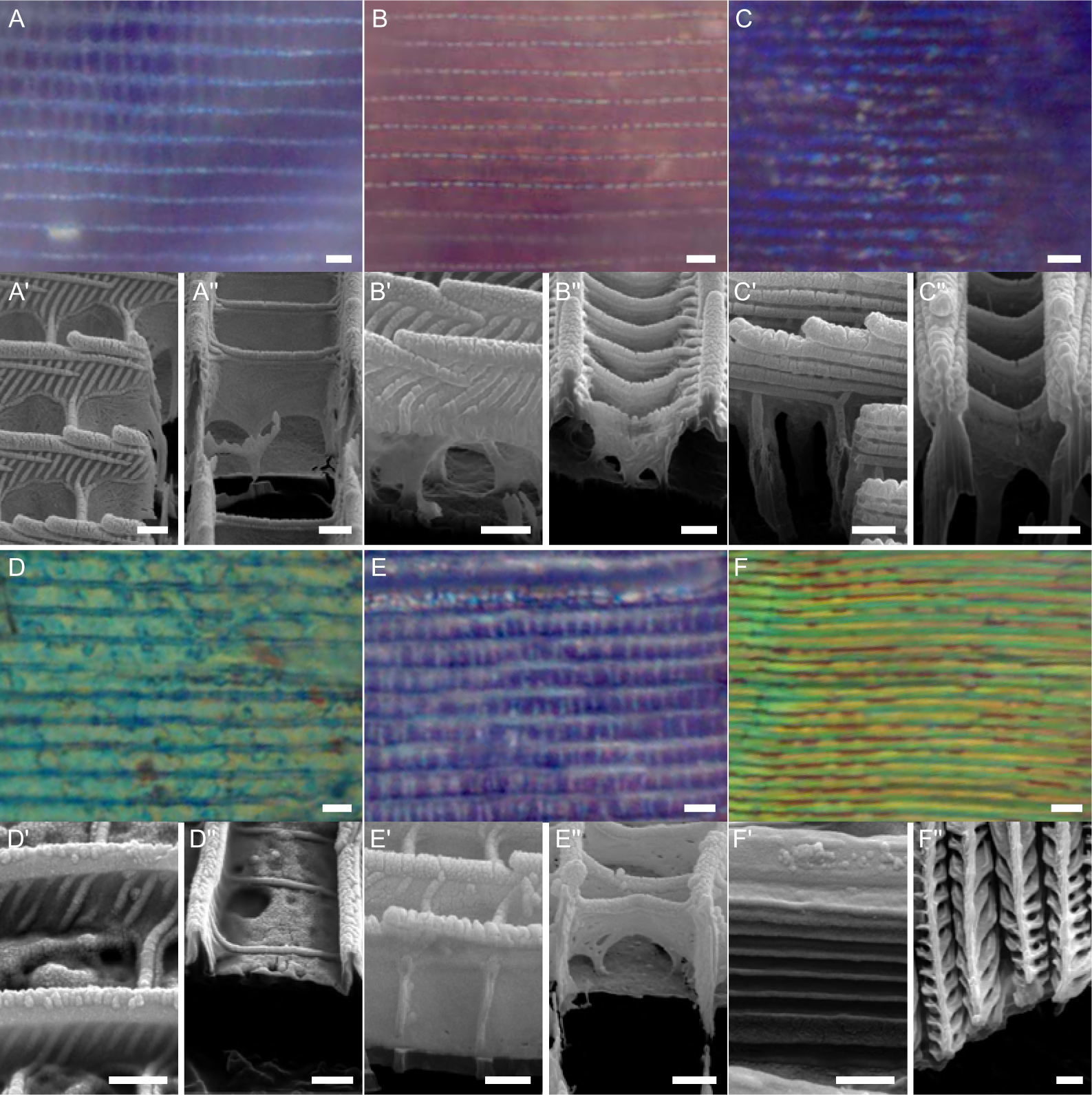
Diversity of ridge and crossrib structural coloration. Optical microscopy images of the abwing side of blue or green cover scales of (A) *Prothoe franck* (Nymphalidae), (B) *Pyrrhopyge hadassa* (Hesperiidae), (C) *Euploea mulciber* (Nymphalidae), (D) *Jalmenus evagoras* (Lycaenidae), (E) *Paralaxita orphna* (Riodinidae), (F) *Trogonoptera brookiana* (Papilionidae). (A’-F’) FIB-SEM images showing sagittal view of scale. (A”-F’’) FIB-SEM images showing transverse view of scale. Scale bars indicate 2 μm on optical microscopy images, and 500 nm on FIB-SEM images.

Our analysis revealed that ridge hue strongly and positively correlated with ridge height, but this correlation was only visible when ridge height was broken down into two separate height intervals (Figure 4A). When the ridges were between 268 and 1036 nm in height, their color hue ranged from about 379 to 594 nm, with a significant positive correlation (Pearson *r*(16)= 0.64 p= 4.5e-03) (Figure 4B). As the ridge height continued to increase, the color sequence began again at cool blues and progressed to warmer hues. This positive correlation was very significant for taller ridges between 1126-2184 nm (Pearson *r*(19)= 0.84 p= 2.1e-06) (Figure 4C).

**Figure 4.**
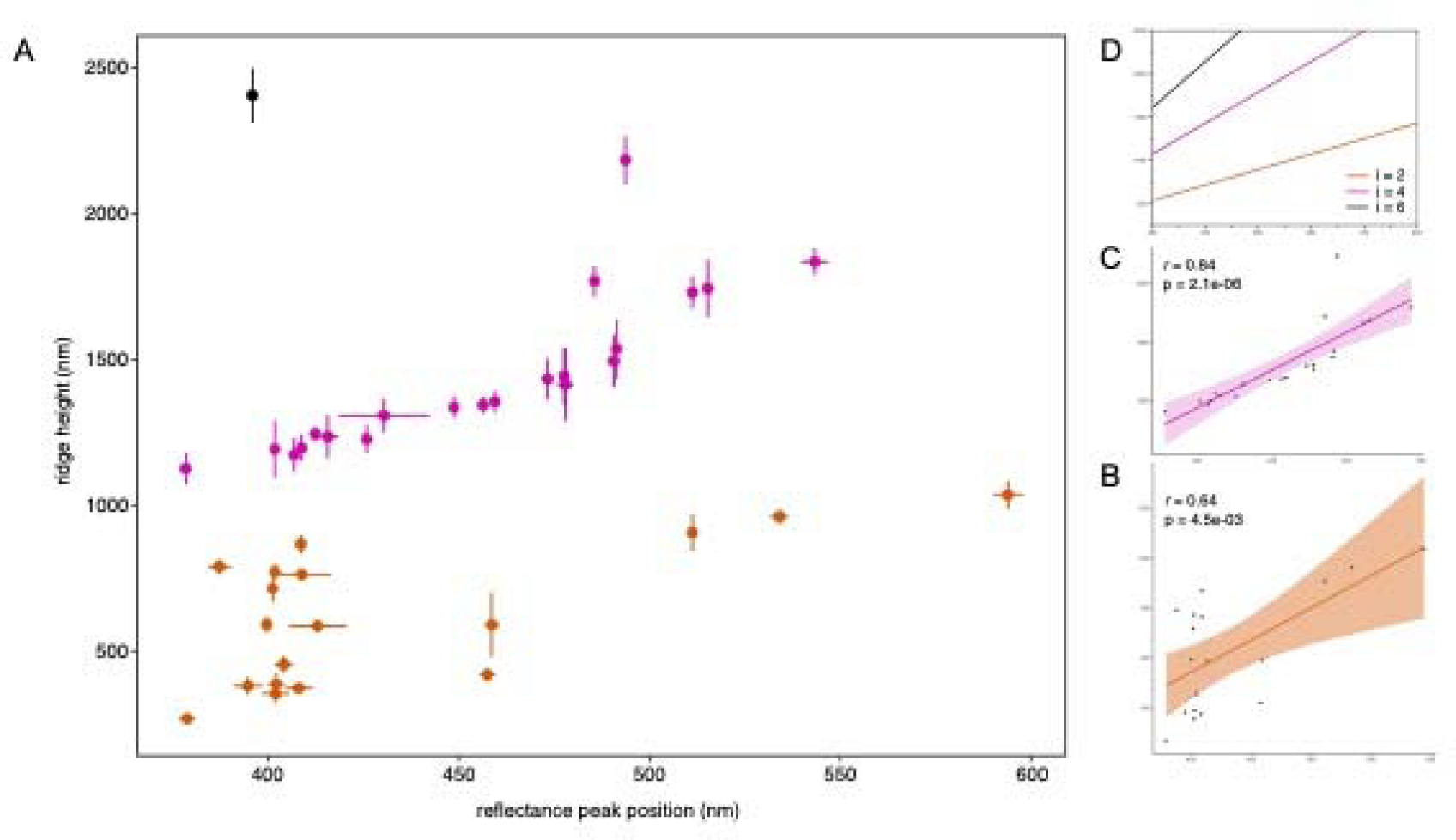
Correlation between ridge height and ridge color hue across species. (A) Scatterplot with error bars showing the whole dataset that includes 40 scales from 35 species (Table S1). Distinct colors were used to help visualize the different cohorts that belong to a different heigh category (step function): 268-1036 nanometers high ridges (orange), 379-594 nanometers high ridges (pink), above 1036 nanometers high ridges (black). (B-C) Linear regression and standard deviation for two sub datasets (= height categories). (D) Plots of peak wavelengths versus the ridge height according to the interference model where *i* equals 2, 4, and 6, respectively.

The correlation between the ridge height and color hue can be explained by a newly described interference model ^30^, in which two light waves propagate through the ridges and through the air next to the ridges (the windows), and interfere with each other when they exit either into the air gap below the ridges, or above the ridges, as light travels back via the same path after reflection from the lower lamina of the scale. When the phase difference between the ridge path and the air path equals an even multiple of Pi (π), the constructive interference of the two waves gives rise to spectral peaks, whose wavelengths/hues (*λ*) are given by:

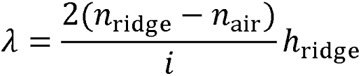

where *i* is an arbitrary even number, *h*_ridge_ is the height of the ridge, and *n*_ridge_ and *n*_air_ are the refractive indices of chitin and air, respectively. The plots where *i* equals 2 and 4 (Figure 4D) represent the cases with the phase difference of 2π and 4π, and they fit the datapoints in Figure 4B and 4C, respectively. This model also predicts a third interval for ridges with the height higher than 1500 nm, which could correspond to our single datapoint in black (Figure 4A).

### Experimental and simulation-based manipulation of ridge height impacts coloration

To go beyond correlations, we manipulated the ridge height in the nymphalid butterfly *Morpho didius* (Figure 5A and 5B). The current model for the metallic blue color of *M. didius* wings assumes that the spatial separation between the stacked lamellae — sometimes referred to as Christmas tree-like reflectors — that protrude from the sides of the ridges is the key structural feature that produces blue color ^8, 31, 32^. The reflected light waves from consecutive lamellae interfere with each other so that the reflected blue wavelengths are intensified via constructive interference, whereas other wavelengths cancel each other out via destructive interference.

**Figure 5.**
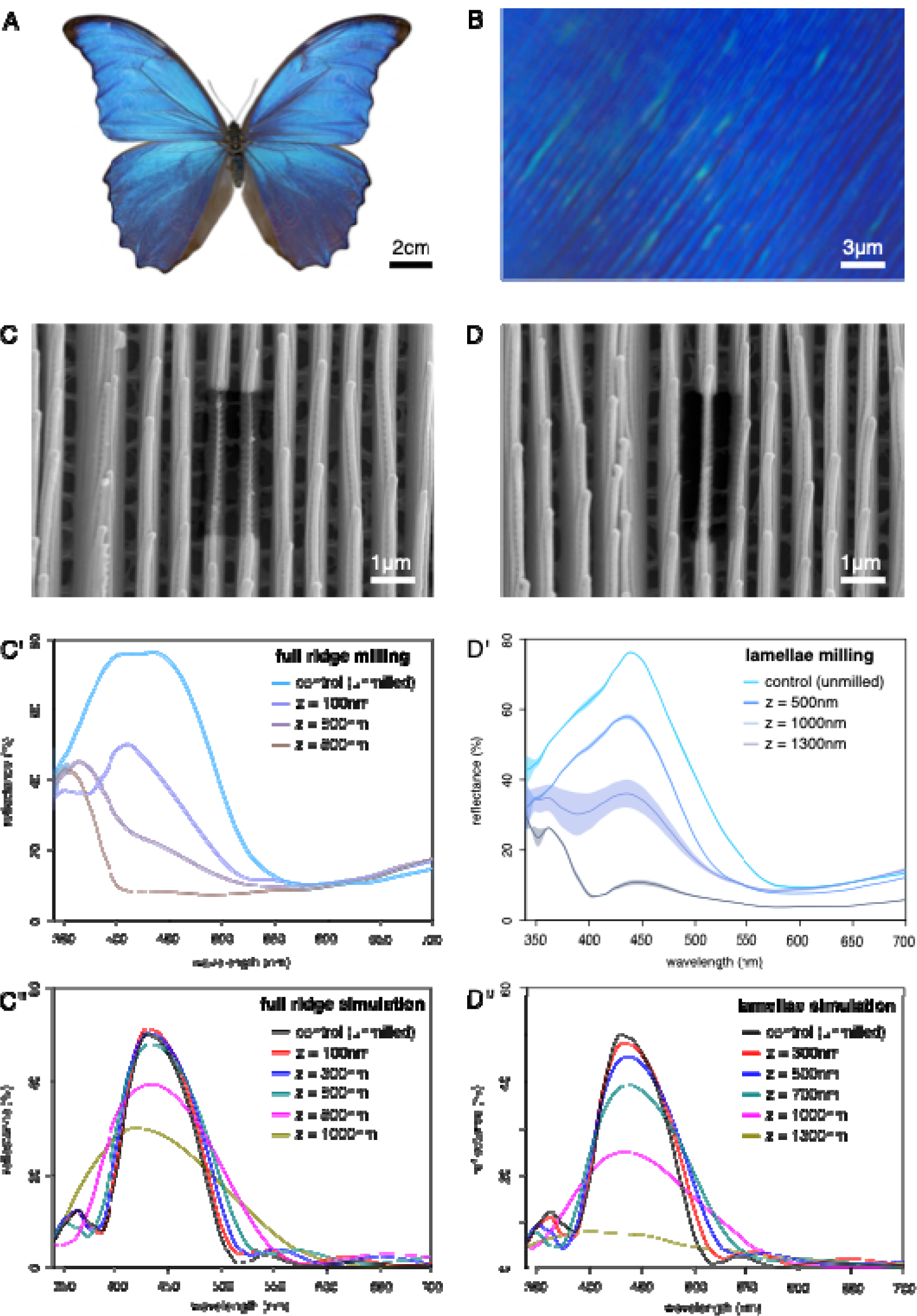
Testing the role of ridge height in *Morpho didius*. (A) Adult specimen of *M. didius* from Peru (photo: Didier Descouens, Muséum d’Histoire Naturelle de Toulouse, CC BY-SA 4.0). (B) Epi-illumination of the wing showing the blue colored ridges (magnification 100X). (C) SEM top view of scale after milling of two consecutive ridges (core + lamellae). (C’) Measured reflectance of intact and milled ridges at different depths. (C’’) Simulated reflectance of intact and milled ridges at different depths. (D) SEM top view of scale after milling of lateral lamellae of the same ridge. (D’) Measured reflectance of intact and milled lamellae at different depths. (D’’) Simulated reflectance of intact and milled lamellae at different depths.

To test the role of ridge height in blue color generation, we performed a series of FIB-SEM millings of the full ridge (core + lamellae) of an uncoated scale at three different depths: 100 nm, 500 nm and 800 nm (Figure 5C), followed by reflectance measurements. We found that the reflection spectrum was UV-shifted when the ridge height was decreased (Figure 5C’). The shift in the hue was also accompanied by a progressive decrease in overall light reflected (Figure 5C’).

To better understand how shortening ridges affects scale hue, we developed an optical model consisting of parallel ridges with lateral lamellae, with dimensions obtained from our SEM images (Figure S3). The simulated reflectance shows a peak around 430 nm that matches the reflection measured for the ridge in real scales (Figure 5D’’). Shortening the ridge height showed a shift towards shorter (UV) wavelengths as the full ridge shortened by 100 to 1000 nm (Figure 5C’’). In summary, both our experimental and *in silico* data show that changes in ridge height alter color hue, and suggests that the height of the central, core part of the ridge alone might control the blue color in *M. didius*.

To disentangle the relative roles of ridge elements further, we selectively removed ridge lamellae from both sides of a single ridge at three different depths (500 nm, 1000 nm and 1300 nm) by FIB milling (Figure 5D). A limitation of gallium ion milling lies in the implantation of ions up to tens of nanometers from the milled area, resulting in local variation in chitin thickness near the edge ^33^. However, our purposely use of low beam voltage and beam current reduced this artifact ^34^. We found that the position of the reflectance peak (∼430 nm) remained unchanged regardless of the depth of milling (Figure 5D’), but light reflection was drastically reduced, being halved by milling to a depth of z = 1000 nm (Figure 5D’). We confirmed these results in our model, by varying the height of the lamellar stack, while keeping constant the height of the core part of the ridge (Figure 5D’’). Similarly, we found that the height of the stack of lamellae is crucial for brilliance and saturation, as previously shown ^31, 32, 35^. In summary, the height of the central, core part of the ridge, and the ridge lamella both seem to be tuned to produce the overall structural blue color of *M. didius*, with the lamellar layers considerably increasing the brightness of the reflected blue light.

## DISCUSSION

Our findings challenge the traditional currently accepted view that thickness variations in the lower lamina of a stereotypical scale is the main ‘evolutionary accessible’ route to alter the scale’s structural color.

First, our study of the artificially selected blue scales of *B. anynana*, showed that ridge and crossrib height and (to a lesser extent), the airgap height beneath these structures, all contributed to the generation of the blue scale hue, previously ascribed solely to changes in lower lamina thickness ^14^.

Second, by revealing a general law between ridge height and color hue, our findings shed new light on a new causative mechanism of structurally colored ridges across nymphalids ^31, 36–39^, pierids ^40, 41^, and lycaenid butterflies ^42^. In many of these species the ridges have a Christmas-tree-like, or Morpho type, structure ^8, 43^, whose prominent side lamella were thought to be the main color-generating features ^8, 31, 32, 35, 44^. However, our comparative analyses, ridge manipulative experiments, and modelling work propose that ridge coloration is produced both by ridge height, and independently from ridge lamella, where the precisely spaced stacked lamellae are primarily amplifying a pre-existing structural hue produced by ridge height.

The optical phenomenon that underlies ridge coloration differs from thin-film interference known to take place in the lower lamina. The structural color coming from the ridges is not due to light interference from reflections produced at the top and bottom of the ridge, as observed in a thin film of chitin. The color results from interference between light which is weakly guided inside the ridge, and light that travels in between consecutive ridges. In other words, light travels slower inside the chitinous ridges compared to light travelling on the outside of the ridges. This phenomenon has already been observed (and modelled) in the lab using non-biological, synthetic nanomaterials. Engineered ridge-like nanostructures made of transparent chitosan (*i.e.*, highly deacetylated chitin) produced the entire structural color palette via variations in height alone ^45^. This is a striking example of how synthetic biology aided our understanding on natural materials, such as butterfly scales.

Interestingly, the ability of simple wall-like ridges and crossribs to produce colors appears to be an ancestral feature of insects and is found in the scales of basal moths and springtails ^19, 46, 47^. Our newfound understanding of how simple wall-like ridges, and the upper lamina in general, is producing color can be harnessed in future to predict the color of fossil insect scales by measuring the dimensions of their scales.

Third, we have shown that the addition of an upper surface can, either increase or decrease a scale’s brilliance, and this may have selective consequences. In cases where a scale’s natural brilliance has been subdued, which often happens in ventral surfaces of butterfly wings ^48, 49^, we hypothesize that this evolved for better crypsis and protection from predators, even if iridescence can sometimes act as camouflage in other insects ^50, 51^. In cases where the upper surface makes scales brighter, as in the case of *Morpho* butterflies ^52^, we hypothesize that this evolved for purposes of sexual signaling ^31^ and/or to confuse predators by signaling unprofitability ^53, 54^.

## MATERIAL AND METHODS

### Butterfly wing scale sampling

*Bicyclus anynana* violet-blue lines were previously generated in our laboratory by artificial selection ^14^. Scales from other butterfly species were sampled on specimens predominantly from the entomological collection of ETH Zurich (ETHZ), the Lee Kong Chian Natural History Museum (LKCNHM) in Singapore, the Museum National d’Histoire Naturelle (MNHN) in Paris, or alternatively from private insect retailers. Details on specimens and sampled scales are provided in Table S1.

### Scanning electron microscopy (SEM)

Scales were mounted on carbon tape, and sputter coated with platinum for 100 s at 40 mA using a JFC-1600 Fine Coat Ion Sputter (JEOL Ltd, Japan). Samples were imaged using a FEI Versa 3D with the following parameters: voltage 10 kV, current 23 pA. Distances between features in scale images were measured using the Line tool implemented in Fiji ^55^. For the ridge-ridge distance, 20 measurements were taken per scale with five scales sampled for each genotype. For the crossrib-crossrib distance, 50 measurements were taken per scale with five scales sampled for each genotype. For the area window, 50 measurements were taken per scale with five scales sampled for each genotype.

### Focused ion beam scanning electron microscopy (FIB-SEM)

Cross sections of wing scales were obtained by FIB milling using the gallium ion beam of the FEI Versa 3D with the following parameters: beam voltage 8 kV, beam current 12 pA, tilt 52°. Milled samples were imaged using a FEI Versa 3D with the following parameters: voltage 10 kV, current 23 pA. The ridge, crossrib, and air gap thicknesses were measured using the Line tool implemented in Fiji ^55^, and corrected for tilted perspective (measured thickness / sin (52°))^56^. Ten independent measurements were taken and averaged.

### Microspectrophotometry

Scales were individually mounted on carbon tape. Reflectance spectra, with a usable range of 340-950 nm, were acquired under normal incidence with a microspectrophotometry set-up including a mercury-xenon light source (Thorlabs, New Jersey, USA) connected to a uSight-2000-Ni microspectrophotometer (Technospex Pte. Ltd, Singapore), using a polished aluminium mirror as a light reference. The microscope’s Nikon TU Plan Fluor objectives have the following specifications: 20x (NA = 0.5), 100x (NA = 0.9). Each measurement was averaged 10 times over an integration time of 100 ms. Reflectance spectra were obtained by averaging three measurements taken at different locations on the scale to account for variability. Reflectance spectra were analysed and plotted using the R package pavo version 2.9 ^57^.

### Color quantification

The reflectance spectra were analyzed to estimate parameters such as wavelength, intensity, and saturation of the reflectance peak. The full-width at half-maximum (FWHM) of the reflectance peak characterizes the desaturation, opposite of saturation ^58^. These parameters were calculated using the peakshape() function of the R package pavo version 2.9 ^57^. The FWHM was calculated using the absolute minimum reflectance of the spectrum. When the R package pavo failed to identify optical peak parameters, we used a split Gaussian function with a Levenberg-Marquardt least square method to fit all the spectral features, using the open-source program Fityk ^59^.

### Optical simulation

The electromagnetic simulations were conducted using a two-dimensional finite-difference time-domain (FDTD) method. A complex refractive index was used for chitin with the refractive index (real part) n = 1.56 and the extinction coefficient (imaginary part) k = 0.06 ^31^. A broadband plane wave was normally incident to periodic arrays of either full ridges, ridges with partially milled core and lamellae, or ridges with partially milled lamellae. The perfectly matched layer-absorbing boundary condition was applied along the light propagation direction to absorb light outside the structure regions. Energy monitors were placed behind the light source to record the reflected light. Input values of the model are indicated in Figure S3. Detailed discussion for the newly described interference model used in Figure 4D is available in ^30^.

### Quantification and statistical analysis

Statistical analysis and plots were done with R 4.2.2 ^60^. The differences in mean height/thickness, distance, or window area between the *B. anynana* wildtype and blue lines were analyzed using a linear mixed model (LME) that allows both fixed and random effects. The rationale was the non-independent, hierarchical nature of the data with multiple measurements taken from each scale, and multiple scales for each individual. LME was run using the nlme 3.1 package ^61^, and different models were compared using the Akaike information criterion (AIC) method. Adjusted *P*-values for different pairwise comparisons were obtained by the Bonferroni post hoc analysis using the multcomp 1.4 package ^62^ and Tukey contrasts. Outputs of the LME tests and adjusted *P*-values for multiple comparisons are shown in Tables S2 and S3.

### Data availability

Measured and simulated reflectance spectra, and measurements of scale geometries have been deposited in Dryad (https://datadryad.org/stash/share/Up0r8AmL-aeoeXaP6POI6uRGfODHdl_68cz0d4LkWDg).

## ACKNOWLEDGEMENTS

We thank Michael Greeff (ETHZ), Rodolphe Rougerie (MNHN) and Wei Song Hwang (LKCHM) for access to entomological collections; Anupama Prakash for help with statistical analyses; Javier Fernandez (SUTD), Joel Yang (SUTD), and Duane Loh (NUS) for discussions, and the Electron Microscopy Facility (EMF, NUS) for use of FIB-SEM. This project was supported by the National Research Foundation (NRF) Singapore, under the Competitive Research Program award NRF-CRP20-2017-0001 to A.M. and V.S. Q.F.R acknowledges the National Natural Science Foundation of China (No. 12304415) and the Natural Science Foundation of Guangdong Province (No. 2023A1515012912).

## AUTHOR CONTRIBUTIONS

C.F. contributed to conceptualization, methodology, investigation, visualization, supervision, writing original draft, draft review and editing. Q.R. contributed to conceptualization, methodology, investigation, writing original draft, draft review and editing. Y.Y.B. contributed to investigation, draft review and editing. V.S. contributed to methodology, draft review and editing. A.M. contributed to conceptualization, methodology, supervision, writing original draft, draft review and editing.

## REFERENCES

1. Srinivasarao, M. Nano-Optics in the Biological World: Beetles, Butterflies, Birds, and Moths. Chem. Rev. 99, 1935–1961 (1999).

2. Vukusic, P. & Sambles, J. R. Photonic structures in biology. Nature 424, 852–855 (2003).

3. Airoldi, C. A., Ferria, J. & Glover, B. J. The cellular and genetic basis of structural colour in plants. Current Opinion in Plant Biology 47, 81–87 (2019).

4. Ghiradella, H. Light and color on the wing: structural colors in butterflies and moths. Appl. Opt. 30, 3492–3500 (1991).

5. Vukusic, P. & Sambles, R. Colour effects in bright butterflies. J. Soc. Dye. Colour. 116, 376–380 (2000).

6. Vukusic, P., Sambles, J. R. & Ghiradella, H. Optical classification of microstructure in butterfly wing-scales. Photonics Sci. News 6, 61–66 (2000).

7. Downey, J. & Allyn, A. Wing-scale morphology and nomenclature. Bull. Allyn Mus. 31, 1–32 (1975).

8. Ghiradella, H. Structure of Iridescent Lepidopteran Scales: Variations on Several Themes. Ann. Entomol. Soc. Am. 77, 637–645 (1984).

9. Ghiradella, H. Structure and Development of Iridescent Lepidopteran Scales: the Papilionidae as a Showcase Family. Ann. Entomol. Soc. Am. 78, (1985).

10. Ghiradella, H. Structure and development of iridescent butterfly scales: Lattices and laminae. J. Morphol. 202, 69–88 (1989).

11. Ghiradella, H. Insect cuticular surface modifications: scales and other structural formations. in Advances in Insect Physiology (eds. Casas, J. & Simpson, S. J.) 38, 135–180 (Elsevier Ltd, 2010).

12. Yeh, P. Optical waves in layered media. (Wiley-Interscience, 2005).

13. Stavenga, D. G., Leertouwer, H. L. & Wilts, B. D. Coloration principles of nymphaline butterflies - Thin films, melanin, ommochromes and wing scale stacking. J. Exp. Biol. 217, 2171–2180 (2014).

14. Wasik, B. R. et al. Artificial selection for structural color on butterfly wings and comparison with natural evolution. Proc. Natl. Acad. Sci. U. S. A. 111, 12109–12114 (2014).

15. Thayer, R., Allen, F. & Patel, N. H. Structural color in Junonia butterflies evolves by tuning scale lamina thickness. Elife 9: e52187, (2020).

16. Bálint, Z. et al. Measuring and Modelling Structural Colours of Euphaedra neophron (Lepidoptera: Nymphalidae) Finely Tuned by Wing Scale Lower Lamina in Various Subspecies. Insects 14, 303 (2023).

17. Stavenga, D. G., Matsushita, A., Arikawa, K., Leertouwer, H. L. & Wilts, B. D. Glass scales on the wing of the swordtail butterfly Graphium sarpedon act as thin film polarizing reflectors. J. Exp. Biol. 215, 657–662 (2012).

18. Finet, C. et al. Multi-scale dissection of wing transparency in the clearwing butterfly Phanus vitreus. J. R. Soc. Interface 20, 20230135 (2023).

19. Zhang, Q. et al. Fossil scales illuminate the early evolution of lepidopterans and structural colors. Sci. Adv. 4, e1700988 (2018).

20. Kilchoer, C., Steiner, U. & Wilts, B. D. Thin-film structural coloration from simple fused scales in moths. Interface Focus 9, 20180044 (2019).

21. Simonsen, T. J. Comparative morphology and evolutionary aspects of the reflective under wing scale-pattern in Fritillary butterflies (Nymphalidae: Argynnini). Zool. Anz. 246, 1–10 (2007).

22. Vukusic, P., Kelly, R. & Hooper, I. A biological sub-micron thickness optical broadband reflector characterized using both light and microwaves. J. R. Soc. Interface 6, S193–S201 (2009).

23. Wilts, B. D., Pirih, P., Arikawa, K. & Stavenga, D. G. Shiny wing scales cause spec(tac)ular camouflage of the angled sunbeam butterfly, Curetis acuta. Biol. J. Linn. Soc. 109, 279–289 (2013).

24. Ren, A. et al. Convergent Evolution of Broadband Reflectors Underlies Metallic Coloration in Butterflies. Front. Ecol. Evol. 8, (2020).

25. Prakash, A., Finet, C., Banerjee, T. Das, Saranathan, V. & Monteiro, A. Antennapedia and optix regulate metallic silver wing scale development and cell shape in Bicyclus anynana butterflies. Cell Rep. 40, 111052 (2022).

26. Mason, C. W. Structural colors in insects. II. J. Phys. Chem. 31, 321–354 (1927).

27. Biró, L. P. et al. Living photonic crystals: Butterfly scales - Nanostructure and optical properties. Mater. Sci. Eng. C 27, 941–946 (2007).

28. Michielsen, K. & Stavenga, D. G. Gyroid cuticular structures in butterfly wing scales: Biological photonic crystals. J. R. Soc. Interface 5, 85–94 (2008).

29. Wilts, B. D., Ijbema, N. & Stavenga, D. G. Pigmentary and photonic coloration mechanisms reveal taxonomic relationships of the Cattlehearts (Lepidoptera: Papilionidae: Parides). BMC Evol. Biol. 14, 160 (2014).

30. Ruan, Q. et al. Reconfiguring Colors of Single Relief Structures by Directional Stretching. Adv. Mater. 34, 2108128 (2022).

31. Vukusic, P., Sambles, J. R., Lawrence, C. R. & Wootton, R. J. Quantified interference and diffraction in single Morpho butterfly scales. Proc. R. Soc. B Biol. Sci. 266, 1403 (1999).

32. Kinoshita, S., Yoshioka, S., Fujii, Y. & Okamoto, N. Photophysics of Structural Color in the Morpho Butterflies. Forma 17, 103–121 (2002).

33. Allen, F. I. et al. Gallium, neon and helium focused ion beam milling of thin films demonstrated for polymeric materials: Study of implantation artifacts. Nanoscale 11, 1403 (2019).

34. Bals, S., Tirry, W., Geurts, R., Zhiqing, Y. & Schryvers, D. High-quality sample preparation by low kV FIB thinning for analytical TEM measurements. Microsc. Microanal. 13, 80–86 (2007).

35. Gralak, B., Tayeb, G. & Enoch, S. Morpho butterflies wings color modeled with lamellar grating theory. Opt. Express 9, 567–578 (2001).

36. Brink, D. J. & Lee, M. E. Confined blue iridescence by a diffracting microstructure: an optical investigation of the Cynandra opis butterfly. Appl. Opt. 38, 5282–5289 (1999).

37. Ćurčić, S. B., et al. Micro- and nanostructures of iridescent wing scales in purple emperor butterflies (Lepidoptera: Apatura ilia and A. iris). Microsc. Res. Tech. 75, 968–976 (2012).

38. Parnell, A. J. et al. Wing scale ultrastructure underlying convergent and divergent iridescent colours in mimetic Heliconius butterflies. J. R. Soc. Interface 15, (2018).

39. Stavenga, D. G., Leertouwer, H. L. & Arikawa, K. Coloration principles of the Great purple emperor butterfly (Sasakia charonda). Zool. Lett. 6, 13 (2020).

40. Ghiradella, H., Aneshansley, D., Eisner, T., Silberglied, R. E. & Hinton, H. E. Ultraviolet reflection of a male butterfly: Interference color caused by thin-layer elaboration of wing scales. Science (80-.). 178, 1214–1217 (1972).

41. Stavenga, D. G., Giraldo, M. A. & Hoenders, B. J. Reflectance and transmittance of light scattering scales stacked on the wings of pierid butterflies. Opt. Express 14, 4880– 4890 (2006).

42. Tilley, R. & Eliot, J. Scale microstructure and its phylogenetic implications in lycaenid butterflies (Lepidoptera, Lycaenidae). Trans. Lepidopterol. Soc. Japan 53, 153–180 (2002).

43. Lippert, W. & Gentil, K. Über lamellare feinstrukturen bei den schillerschuppen der schmetterlinge vom urania- und morpho-typ. Zeitschrift für Morphol. und Ökologie der Tiere 48, 115–122 (1959).

44. Tabata, H., Kumazawa, K., Funakawa, M., Takimoto, J. I. & Akimoto, M. Microstructures and optical properties of scales of butterfly wings. Opt. Rev. 3, 139– 145 (1996).

45. Raut, H. K. et al. The Height of Chitinous Ridges Alone Produces the Entire Structural Color Palette. Adv. Mater. Interfaces 9, 2201419 (2022).

46. D’Alba, L., Wang, B., Vanthournout, B. & Shawkey, M. D. The golden age of arthropods: Ancient mechanisms of colour production in body scales. J. R. Soc. Interface 16, 20190366 (2019).

47. Vanthournout, B. et al. Springtail coloration at a finer scale: Mechanisms behind vibrant collembolan metallic colours. J. R. Soc. Interface 18, 20210188 (2021).

48. Oliver, J. C., Robertson, K. A. & Monteiro, A. Accommodating natural and sexual selection in butterfly wing pattern evolution. Proc. R. Soc. B Biol. Sci. 276, 2369–2375 (2009).

49. Allen, C. E., Zwaan, B. J. & Brakefield, P. M. Evolution of sexual dimorphism in the lepidoptera. Annu. Rev. Entomol. 56, 445–464 (2011).

50. Kjernsmo, K. et al. Iridescence impairs object recognition in bumblebees. Sci. Rep. 8, (2018).

51. Kjernsmo, K. et al. Iridescence as Camouflage. Curr. Biol. 30, 551–555 (2020).

52. Giraldo, M. A. & Stavenga, D. G. Brilliant iridescence of Morpho butterfly wing scales is due to both a thin film lower lamina and a multilayered upper lamina. J. Comp. Physiol. A Neuroethol. Sensory, Neural, Behav. Physiol. 202, (2016).

53. Pinheiro, C. E. G., Freitas, A. V. L., Campos, V. C., DeVries, P. J. & Penz, C. M. Both Palatable and Unpalatable Butterflies Use Bright Colors to Signal Difficulty of Capture to Predators. Neotropical Entomology 45, 107–113 (2016).

54. Le Roy, C. et al. Adaptive evolution of flight in Morpho butterflies. Science (80-.). 374, 1158–1162 (2021).

55. Schindelin, J., et al. Fiji: An open-source platform for biological-image analysis. Nature Methods 9, (2012).

56. Villinger, C. et al. FIB/SEM tomography with TEM-like resolution for 3D imaging of high-pressure frozen cells. Histochem. Cell Biol. 138, 549–556 (2012).

57. Maia, R., Gruson, H., Endler, J. A. & White, T. E. pavo 2: New tools for the spectral and spatial analysis of colour in r. Methods Ecol. Evol. 10, (2019).

58. Oppenheim, A. V., Willsky, A. S. & Nawab, S. H. Signals & systems. (Prentice-Hall, Inc., 1997).

59. Wojdyr, M. Fityk: A general-purpose peak fitting program. J. Appl. Crystallogr. 43, 1126–1128 (2010).

60. R Core Team. R: a language and environment for statistical computing. (2021).

61. Pinheiro, J., Bates, D. & R Core Team. nlme: linear and nonlinear mixed effects models. R package version 3.1–162. (2023).

62. Hothorn, T., Bretz, F. & Westfall, P. Simultaneous inference in general parametric models. Biometrical Journal 50, 346–363 (2008).

63. Espeland, M. et al. A Comprehensive and Dated Phylogenomic Analysis of Butterflies. Curr. Biol. 28, 770–778 (2018).

