## Supplementary material for "Ridge and crossrib height of butterfly wing scales is a toolbox for structural color diversity": Figure S1

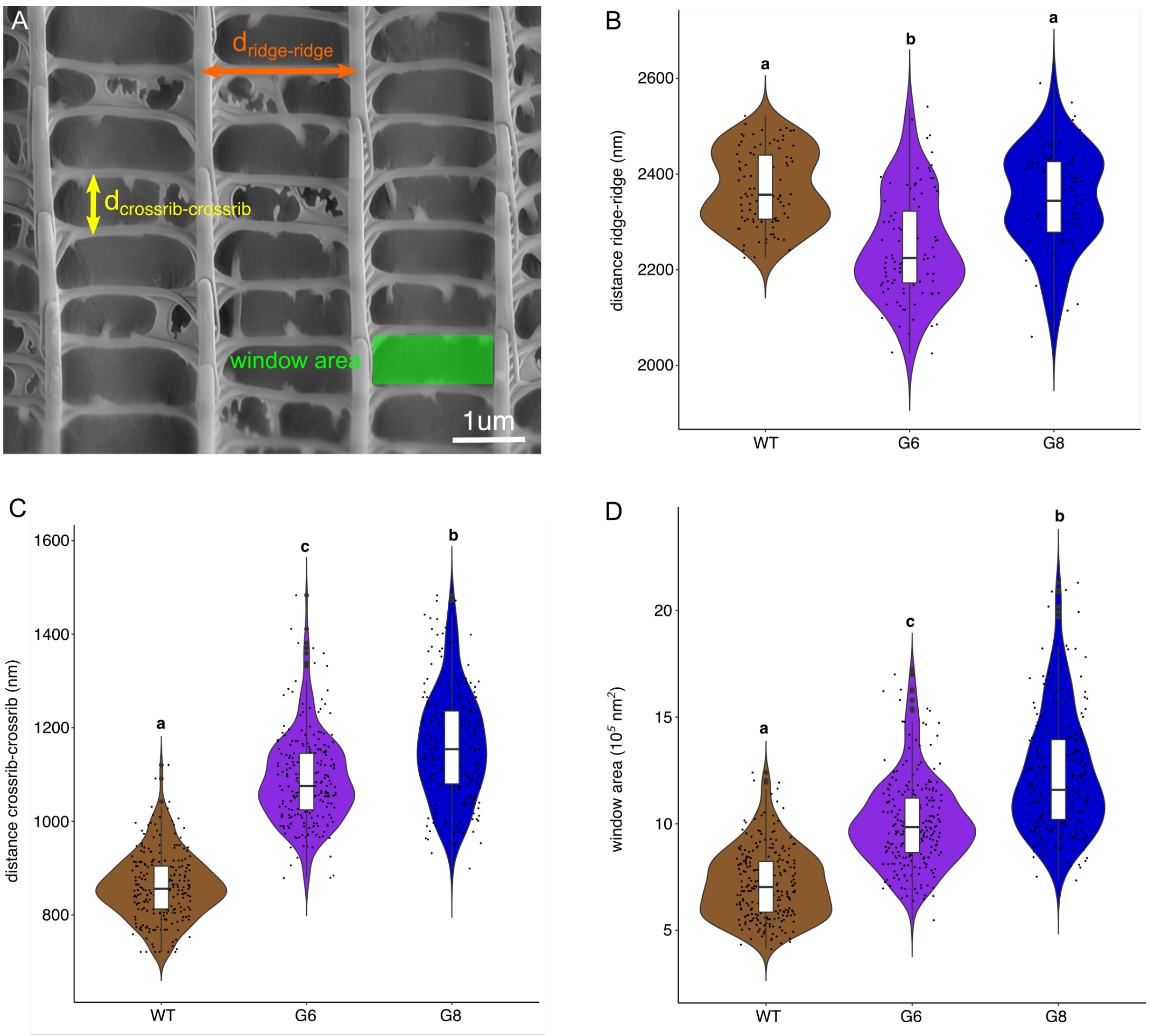


**Figure S1.** Comparison of the distance ridge-ridge, distance cross-cross-rib, and window area between WT, generation G6, and generation G8 individuals in *B. anynana*. (A) Schematic of the measured geometries. (B) The distance between ridges does not seem to follow a trend over selection. (C) The distance between crossribs significantly increases over generations (see tables S2 and S3 for statistics). (D) The change in distances led to a significant enlargement of the window area (tables S2 and S3), which makes the colour generated by the lower lamina more visible.
