## Supplementary material for "Ridge and crossrib height of butterfly wing scales is a toolbox for structural color diversity": Figure S2

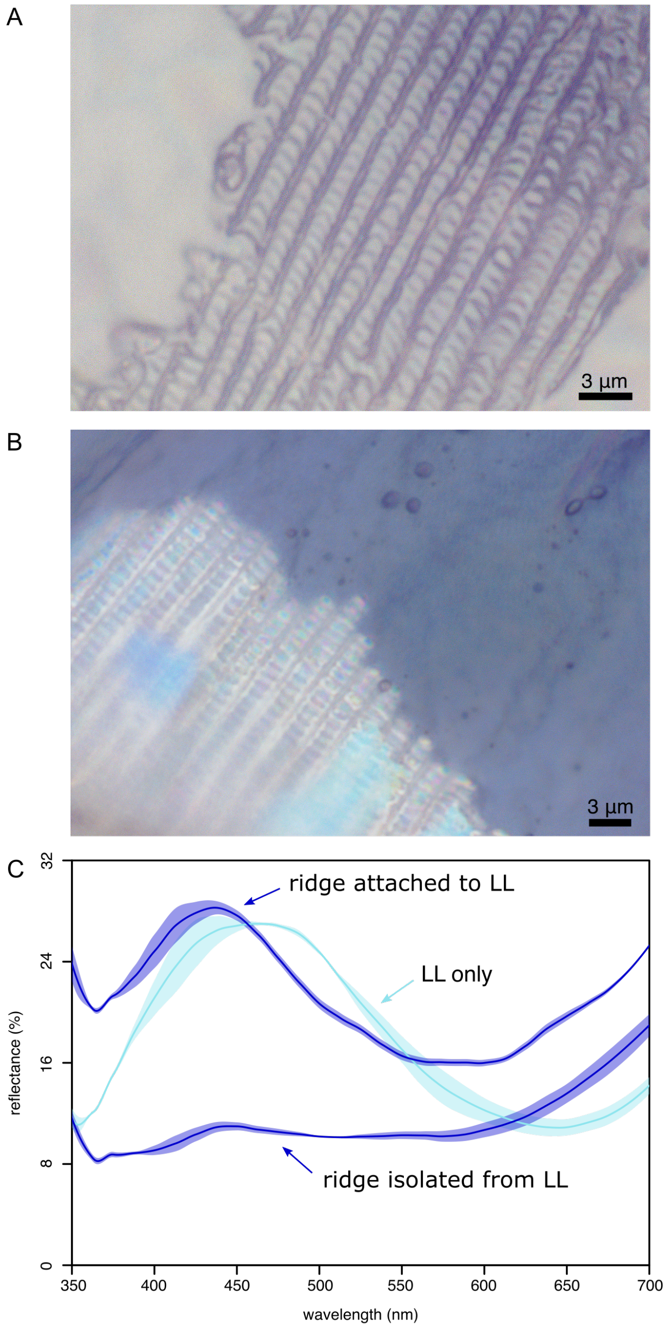


**Figure S2.** Comparison of ridge reflectance between upper surface on intact scale and upper surface isolated from lower lamina. A ground scale from a selected *B. anynana* individual (G8) was isolated and its upper surface was partly removed, leaving the lower lamina (LL) exposed. Then, the reflectance spectrum of ridges was measured. (A) Isolated upper surface mounted on glass side. (B) Scale after partial removal of the upper surface: the lower lamina is visible on the right part of the scale. (C) The reflectance peak of the ridge is located at the same position regardless of the methodology, *i.e.*, the upper surface isolated or not from the lower lamina. Note: the intensity of the ridge reflectance is higher on the intact scale upper because the lower lamina prevents the back scattering of white light. In addition, the reflectance of the abwing (upperside) lower lamina after upper surface removal is shown in cyan. The difference in reflectance peaks confirms that our spectrophotometry set up is able to measure the reflectance of an individual ridge, without or very little contamination from the underneath lower lamina.
