## Supplementary material for "Ridge and crossrib height of butterfly wing scales is a toolbox for structural color diversity": Figure S3

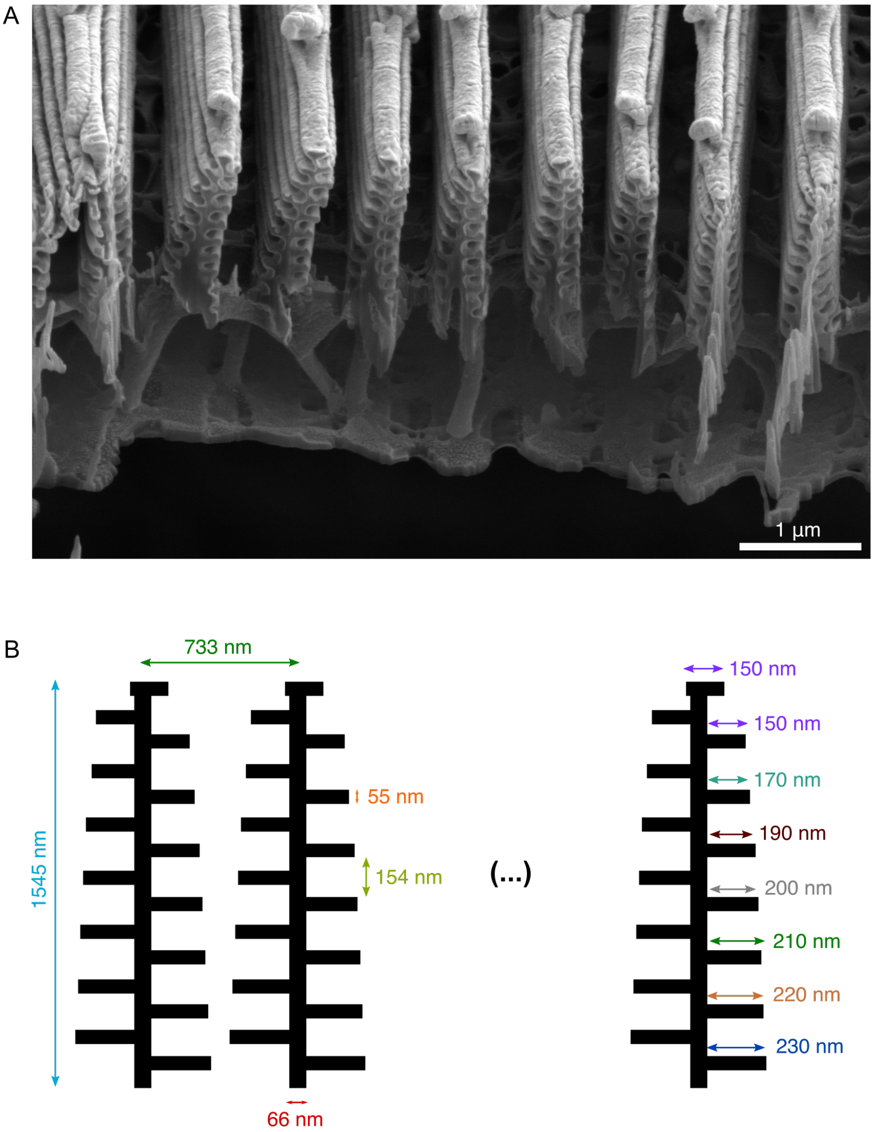


**Figure S3.** Specifications of the *Morpho didius* model. (A) FIB-SEM cross-section used to measure distances and geometries. (B) Parameter values used as input for the simulation of *M. didius* ridges and lamellae.
