## Supplementary material for "Ridge and crossrib height of butterfly wing scales is a toolbox for structural color diversity": Tables S1&S2

**Table S1.** Taxon sampling. ETHZ = entomological collection of ETH Zurich, LKCNHM = Lee Kong Chian Natural History Museum (Singapore), MNHN = Museum National d'Histoire Naturelle (Paris), * The Bugmaniac Insect Shop, ** Stratford-upon-Avon Butterfly Farm, *** MARL Insect and Butterfly Culture, D = dorsal, V = ventral, FW = forewing, HW = hindwing.

| Family | Subfamily | Tribe | Genus | Species | Source | Specimen details | Sex | Sampled scales |
|---|---|---|---|---|---|---|---|---|
| Hesperiidae | Hesperiinae | Moncini | *Vettius* | *phyllus* | ETHZ | drawer 333 | M | DFW blue |
| Lycaenidae | Miletinae | Miletini | Allotinus | subviolaceus | LKCNHM | ZRC.7.04178 (Malaysia) | M | DFW blue |
| | Theclinae | Zesiusini | *Jalmenus* | *evagoras* | ETHZ | drawer 322 (New Holland) | | DFW cyan |
| Nymphalidae | Danainae | Danaini | *Amauris* | *niavius* | ETHZ | drawer 130-135 | | VFW silver |
| | Charaxinae | Anaeomorphini | *Anaeomorpha* | *splendida* | ETHZ | drawer 166 (Colombia, 1931) | M | DFW blue green |
| | Charaxinae | Charaxini | *Charaxes* | *numenes* | ETHZ | drawer 153-161, 165 (Cameroon) | M | DHW blue |
| | Satyrinae | Haeterini | *Cithaerias* | *esmeralda* | ETHZ | drawer 291 (Amazonia, 1921) - labelled *Callitaera esmeralda* | F | DHW blue |
| | Limenitidinae | Limenitidini | *Cymothoe* | *aemilius* | ETHZ | drawer 234-235 (Cameroon, 1925) | M | DFW + VFW cyan |
| | Cyrestinae | Cyrestini | *Cyrestis* | *achates* | eBay | Indonesia | M | VFW blue |
| | Pseudergolinae | | *Dichorragia* | *nesimachus* | ETHZ | drawer 151 | | VFW blue |
| | Apaturinae | | *Doxocopa* | *laurentia* | ETHZ | drawer 152 - labelled as *Doxocopa seraphina* | M | DFW cyan + violet |
| | Biblidinae | Dynaminini | *Dynamine* | *mylitta* | ETHZ | drawer 239-240 | M | DHW green + VHW blue |
| | Satyrinae | Elymniini | *Elymnias* | *malelas* | ETHZ | drawer 298-299 | M | DFW blue |
| | Biblidinae | Epiphilini | *Epiphile* | *kalbreyeri* | ETHZ | drawer 212-214 (Colombia, 1924) | M | DHW blue |
| | Limenitidinae | Adoliadini | *Euphaedra* | *cyparissa* | ETHZ | drawer 236-238 (Sierra Leone) | M | DHW or FHW green |
| | Nymphalinae | Coeini | *Historis* | *odius* | ETHZ | drawer 241 | M | VFW violet |
| | Nymphalinae | Junoniini | *Junonia* | *orithya* | field | Coney Island, Singapore (Permit No: NP/RP 14-063-2) | M | DHW blue |
| | Heliconiinae | Vagrantini | *Lachnoptera* | *iole* | * | Central African Republic | | VHW blue |
| | Satyrinae | Satyrini | *Lethe* | *sinorix* | LKCNHM | ZRC.7.03817 (Malaysia, 1940) | | VHW purple |
| | Limenitidinae | Limenitidini | *Limenitis* | *astyanax* | ETHZ | drawer 232-233 | | DHW + VHW green |
| | Cyrestinae | Cyrestini | *Marpesia* | *corinna* | * | Peru | M | DHW violet |
| | Satyrinae | Melanitini | *Melanitis* | *leda* | LKCNHM | ZRC.7.03865 (Malaysia, 1921) | | VHW blue eyespot |
| | Satyrinae | Morphini | *Morpho* | *anaxibia* | ETHZ | drawer 258-278 (Brazil) | M | DHW blue |
| | Satyrinae | Morphini | *Morpho* | *didius* | eBay | Peru | M | ground + cover DHW blue |
| | Satyrinae | Morphini | *Morpho* | *peleides* | ** | | M | ground + cover DHW blue |
| | Satyrinae | Morphini | *Morpho* | *sulkowskyi* | ETHZ | drawer 258-278 (Nouvelle-Grenade, 1884) | M | DHW cyan |
| | Nymphalinae | Victorinini | *Napeocles* | *jucunda* | ETHZ | drawer 203 (Amazonia) | M | DFW cyan |
| | Charaxinae | Pallini | *Palla* | *violinitens* | ETHZ | drawer 162 (Cameroon) | M | VFW violet |
| | Biblidinae | Ageroniini | *Panacea* | *procilla* | ETHZ | drawer 245 (Peru) | | DFW cyan green |
| | Limenitidinae | Parthenini | *Parthenos* | *sylvia* | *** | The Philippines | M | DHW violet |
| | Charaxinae | Preponini | *Prepona* | *pheridamas* | ETHZ | drawer 166-182 | | DFW cyan |
| | Charaxinae | Prothoini | *Prothoe* | *franck* | MNHN | | M | DFW purple |
| Papilionidae | Papilioninae | Papilionini | *Chilasa* | *paradoxa* | LKCNHM | ZRC_ENT00024516 (Malaysia, 1965) - labelled as *Chilasa paradoxa aenigma* | F | DFW pale blue |
| Riodinidae | Riodininae | Eurybiini | *Alesa* | *prema* | ETHZ | drawer 312 (Amazonia) | M | DHW green |
| | Nemeobiinae | Nemeobiini | *Paralaxita* | *orphna* | LKCNHM | ZRC.7.02137 (Siam, 1922) - labelled as *Laxita orphna laocoon* | M | VHW blue |

**Table S2.** LME test results for the comparison of geometries between WT and artificially selected blue ground scales in *B. anynana* (see relevant plots in Figure 2).

| factor | numDF | denDF | F-value | P-value |
|---|---|---|---|---|
| *ridge height* | | | | |
| genotype | 2 | 147 | 26,25 | <0.0001 |
| *crossrib height* | | | | |
| genotype | 2 | 147 | 69,24 | <0.0001 |
| *sub ridge air thickness* | | | | |
| genotype | 2 | 143 | 86,2 | <0.0001 |
| *sub crossrib air thickness* | | | | |
| genotype | 2 | 147 | 35,97 | <0.0001 |
| *ridge + air height* | | | | |
| genotype | 2 | 143 | 115,12 | <0.0001 |
| *crossrib + air height* | | | | |
| genotype | 2 | 147 | 78,06 | <0.0001 |
| *ridge-ridge distance* | | | | |
| genotype | 2 | 293 | 48,41 | <0.0001 |
| *crossrib-crossrib distance* | | | | |
| genotype | 2 | 743 | 835,74 | <0.0001 |
| *window area* | | | | |
| genotype | 2 | 743 | 378,52 | <0.0001 |
