## Supplementary material for "Ridge and crossrib height of butterfly wing scales is a toolbox for structural color diversity": Table S3

**Table S3.** Statistical test results for the multiple comparisons of geometries among different genotypes obtained from Tukey post-hoc tests (see relevant plots in Figure 2).

| comparison | estimate | standard error | z-value | Pr(>\|z\|) |
|---|---|---|---|---|
| *ridge height* | | | | |
| G8 - G6 == 0 | 41,29 | 7,14 | 5,78 | <1e-05 *** |
| WT - G6 == 0 | -6,35 | 7,14 | -0,89 | 0,65 |
| WT - G8 == 0 | -47,64 | 7,14 | -6,67 | <1e-05 *** |
| *crossrib height* | | | | |
| G8 - G6 == 0 | 48,2 | 5,75 | 8,38 | <1e-04 *** |
| WT - G6 == 0 | -14,16 | 4,92 | -2,88 | 0.01 * |
| WT - G8 == 0 | -62,36 | 5,35 | -11,66 | <1e-04 *** |
| *sub ridge air thickness* | | | | |
| G8 - G6 == 0 | 69,52 | 10,43 | 6,66 | 8.08e-11 *** |
| WT - G6 == 0 | -67,48 | 10,43 | -6,47 | 2.99e-10 *** |
| WT - G8 == 0 | -137,01 | 10,43 | -13,13 | <2e-16 *** |
| *sub crossrib air thickness* | | | | |
| G8 - G6 == 0 | 72,71 | 11,76 | 6,18 | 1.85e-09 *** |
| WT - G6 == 0 | -29,2 | 8,47 | -3,45 | 0.0017 ** |
| WT - G8 == 0 | -101,91 | 12,02 | -8,48 | <2e-16 *** |
| *ridge + air height* | | | | |
| G8 - G6 == 0 | 110,81 | 12,25 | 9,05 | <2e-16 *** |
| WT - G6 == 0 | -73,84 | 12,25 | -6,03 | 4.99e-09 *** |
| WT - G8 == 0 | -184,65 | 12,25 | -15,07 | <2e-16 *** |
| *crossrib + air height* | | | | |
| G8 - G6 == 0 | 120,91 | 13,33 | 9,07 | <2e-16 *** |
| WT - G6 == 0 | -43,36 | 10,11 | -4,29 | 5.39e-05 *** |
| WT - G8 == 0 | -164,27 | 13,15 | -12,49 | <2e-16 *** |
| *ridge-ridge distance* | | | | |
| G8 - G6 == 0 | 97,54 | 14,66 | 6,65 | 8.59e-11 *** |
| WT - G6 == 0 | 122,19 | 12,6 | 9,7 | <2e-16 *** |
| WT - G8 == 0 | 24,65 | 13,17 | 1,87 | 0,184 |
| *crossrib-crossrib distance* | | | | |
| G8 - G6 == 0 | 70. 70 | 9,59 | 7,37 | 4.99e-13 *** |
| WT - G6 == 0 | -227,2 | 7,52 | -30,21 | <2e-16 *** |
| WT - G8 == 0 | -297,89 | 8,49 | -35,07 | <2e-16 *** |
| *window area* | | | | |
| G8 - G6 == 0 | 224425 | 24137 | 9,3 | <2e-16 *** |
| WT - G6 == 0 | -289239 | 16181 | -17,87 | <2e-16 *** |
| WT - G8 == 0 | -513664 | 21378 | -24,03 | <2e-16 *** |
